# The choice-wide behavioral association study: data-driven identification of interpretable behavioral components

**DOI:** 10.1101/2024.02.26.582115

**Authors:** David B. Kastner, Nicole O. Yokota, Christina Y. Lee, Cristofer Holobetz, Ali Chaudhry, Greer Williams, Jane Ton, Mihika Mehra, Viktor Kharazia, Dilworth Y. Parkinson, Joseph P. Romano, Peter Dayan

**Author notes:** Correspondence:* David B. Kastner.

## Abstract

Behavior contains rich structure across many timescales. However, there is a dearth of methods to identify relevant components, especially over the longer periods required for learning and decision-making. Inspired by the goals and techniques of genome-wide association studies, we present a data-driven method—the choice-wide behavioral association study: CBAS—that systematically identifies such behavioral features. CBAS extracts sequences of actions or choices that either significantly differ between groups or correlate with a covariate of interest, using a powerful resampling-based method to correct for multiple comparisons. We illustrate CBAS through application to tasks performed by flies, rats and humans, showing that it provides unexpected and interpretable information in each case.

## INTRODUCTION

Understanding how behavior differs between or among groups of humans or other animals is critical for generating and testing hypotheses about the functional role of genes, neural circuits, and regions of the brain. It is also essential for characterizing neurologic and psychiatric dysfunction^1^. However, behavior is highly complex, evolving over multiple timescales and exhibiting substantial path dependencies due to individual experience^2–6^. To deal with these complexities, it is increasingly popular to collect large datasets through automated behavioral paradigms, and to analyze them using computational and machine learning methods^7–16^. There are two main computational approaches^17,18^: top-down/model-dependent, and bottom-up/data-driven. Model-dependent methods are substantially more prevalent in the analysis of behavior; however, data-driven methods dominate in other areas with rich and complex findings and afford the possibility of addressing some of the many limitations of top-down approaches.

In model-dependent analyses, behavioral data, such as choices in a decision-making task, are processed under the specific assumptions of a hypothesis or model. If the model or hypothesis is correct, this is highly efficient, since large volumes of data can be reduced to a handful of semantically meaningful parameters, such as learning rates or differential sensitivity to rewards or punishments. By finding specific parameters whose values differ significantly between groups, or correlate with facets of those groups, model-dependent analyses conflate identification with interpretation. That is both their strength and their weakness. If the model is correct, it allows for the difference or correlation to be directly interpreted in terms of the parameters concerned; however, if the model is incorrect or limited, confirmation bias^19^imposes an incorrect interpretation on the difference or correlation. That is, even when substantial effort is put into building multiple alternative models, and comparing them in a statistically rigorous manner, it remains possible that the best fitting model nevertheless fails to characterize the behavior properly. This limits the interpretability of the model or the interrogation of hypotheses. In the worst cases, model and hypothesis-based analyses offer post-hoc explanations for any difference that can be found in a behavioral dataset^20^.

Data-driven analyses start from the other end, characterizing behavior without relying on parametric assumptions, and making very few assumptions about how the data are generated. Some of these approaches are unsupervised, for instance finding clusters in behavioral space and comparing them between groups^12,21,22^; others are more supervised, directly looking for discriminative differences between populations or individuals^7–10^. By not trying to shoehorn data into a limited set of parameters, these methods should be more sensitive; however, comparisons between groups pose severe statistical challenges, because of the complexity and dimensionality of behavioral datasets; and interpretation remains a challenge.

Modern machine learning methods^23–25^ can sit between model-dependent and data-driven camps. They have the particular benefit of providing useful lower bounds for how well hypothesis-driven models should fit but often struggle with convincing interpretability and data efficiency.

Here, we present the choice-wide behavioral association study (CBAS), a novel data-driven analysis method designed to identify sequences of choices (or other discrete behavioral features) that differ between or among groups. CBAS has two components: 1) breaking down behavior into a comprehensive language that allows comparisons to be made between two groups or to assess correlation with a covariate of interest; and 2) using rigorous, resampling-based, corrections to account for the resulting large number of comparisons and to maintain statistical power despite statistical complexity in the data. CBAS breaks the conflation between detection and interpretation of model-dependent analyses because it is designed simply to identify differences or find correlations with a covariate. The presence (or absence) of that significance is interpretation-free. However, by design, the output of CBAS provides a rich and rigorous substrate for interpreting behavior, for testing the adequacy of models and hypotheses concerning the source of behavioral differences, and our method provides a general data-driven framework to advance our resulting understanding.

## RESULTS

### Choice as a common discretization for behavior

The development of CBAS was motivated by the powerful data-driven approaches of genome-wide association studies (GWAS), whole exome sequencing (WES) and whole genome sequencing (WGS)^26^. Prior to the wide-scale implementation of these methods, candidate gene studies that attempted to associate candidate genes with phenotypes were often underpowered and failed to replicate^27–29^. It has been argued that many data-driven behavioral analyses suffer the same problems^30^. GWAS/WES/WGS famously solved the problem of statistical reliability by employing rigorous multiple-comparisons correction. These corrections depend on the discrete representation afforded by base-pairs^31,32^. CBAS follows an equivalent statistical path, and also requires a similarly appropriate, and potentially general, discretization, but for behavior. In many cases, choices provide just such a discretization (Fig 1a).

**Figure 1.**
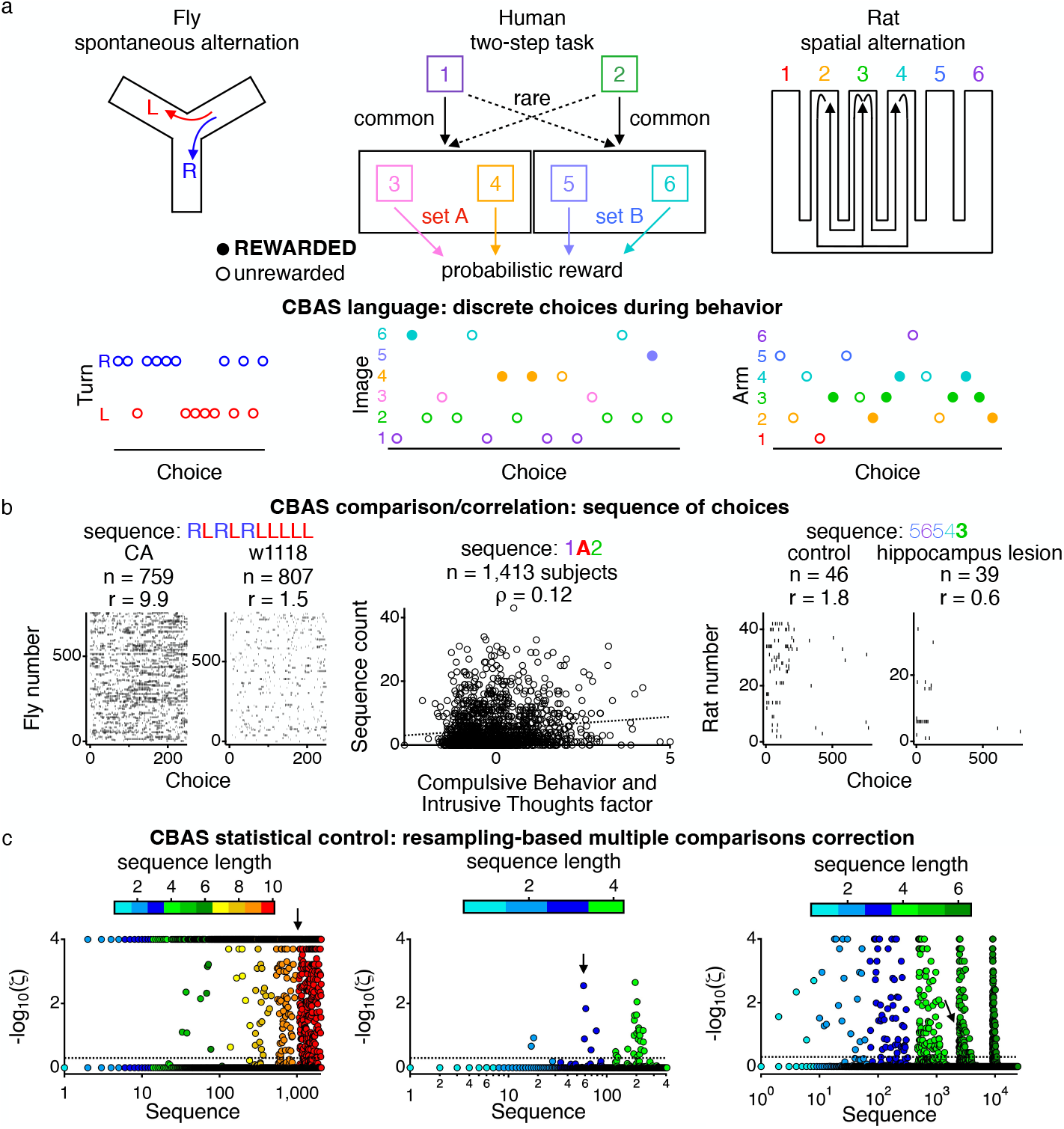
Choice-Wide Behavioral Association Study (CBAS) is applicable across diverse tasks and species. (**a** The starting point for the three behavioral tasks we consider is to decompose the actions of the subjects into a series of discrete choices. Left: Fly y-maze (top), and example snippet of the choices of left (red) and right (blue) turns (bottom) of an individual fly. Data from Buchanan et al.^31^. Middle: Two-step task, performed by human subjects (top), and an example snippet of choices of an individual subject (bottom). The colors correspond to the different images chosen (bottom). Data from Gillan et al.^43^. Right: Spatial alternation behavior, performed by rats (top), and an example snippet of choices of an individual rat (bottom). Colors correspond to the arm chosen by the rat. The rats are rewarded if they visit arms 2, 3, and 4 by alternating visits to arms 2 and 4 after visiting arm 3 (arrows). The flies were not rewarded for their choices; for the middle and right panels, filled circles indicate that reward was received. (**b**) CBAS breaks down the task into all sequences of choices up to a defined total length and compares the usage rate of the sequences between two groups (flies and rats) or correlates the usage rate of each sequence with a covariate of interest (humans). In all cases, CBAS controls statistically for the large number of sequence comparisons. Left: The occurrence of a single sequence of 10 turns in the two fly strains during the first 250 turns of the spontaneous alternation behavior. The rows show the occurrence of that sequence of turns for all individual flies from each strain. For this example sequence, the CA strain has a rate of sequence usage of 9.9 per animal. The w1118 strain has a rate of sequence usage of just 1.5 per animal. This difference in rate is significant with *ζ*= 1*x*10^−4^, corrected for the multiple comparisons. Middle: Correlation between the participants’ CBIT scores and their usage of the particular sequence 1**A**2, which means choosing image 1, then making a choice in set A and getting rewarded and then choosing image 2. The usage of this sequence shows a correlation of 0.12 (*ζ*< _2_.8*x*10^−3^) with the CBIT score of each participant. Dotted line shows the linear fit to the data. Right: The occurrence of a single sequence of arm choices 5654**3** (unrewarded for the first 5 choices but then rewarded for the move into the central arm of the rules, arm 3) in the two groups of rats (control and hippocampal lesioned) during the first 800 arm choices. The rows show the occurrence of that sequence of choices for all individual rats from each group. Control rats use this sequence at a rate of sequence usage of 1.8 per animal. Hippocampal lesioned rats use this sequence at a rate of sequence usage of 0.6 per animal. This difference in rate is significant with *ζ*= 0.047. (**c**) CBAS Manhattan plot displays the p-value for each sequence evaluated in the 3 datasets. The sequences are ordered based on the number of choices each involves (shown by colors, up to the total length considered), and they are displayed on a log scale to make all sequence lengths visible. Within a given sequence length, the sequences are ordered based on frequency of occurrence in the entire dataset. The horizontal dotted line indicates the significance threshold of *γ*=5% control of the median false discovery proportion (*ζ*= 0.5). A total of 10,000 resamples were used to calculate the p-values, making the maximum value on the plot 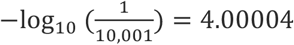 The location of the example sequences shown in **b** are indicated by arrows.

We apply CBAS to three tasks in three diverse species—flies, humans, and rats—to show the breadth of applicability of CBAS. For simplicity and clarity, we describe the motivation and development of CBAS using one of the tasks and then show its application to the other tasks below. The task that we will initially use to describe CBAS involves drosophila walking on a y-maze (Fig 1a; left). The flies are tracked as they spontaneously make the choice of going to either the left or right arm after leaving a current arm^31^. This left/right discretization of the task has enabled many conclusions to be drawn about the genetic nature of individual variability^31– 36^. Here, we compare the choices of two outbred strains of fly, Cambridge-A (CA) and w1118, from a publicly available dataset^31^. The w1118 strain is the background strain for many transgenic flies^31^. Analyses in the original paper^31^ hypothesized a potential source of the difference between the groups: handedness, or the preference for left or right turns, by the individual flies; however, there are many different ways in which that might manifest, and the rich dataset combined with CBAS allows multiple possibilities to be evaluated.

### CBAS evaluates all sequences of choices up to a set length

Using choice as the basis of the comparison for the behavioral analysis requires an additional consideration beyond what is done for the genome with base-pairs. For GWAS/WES/WGS, an individual base pair can be a meaningful unit of information (although this is only a partial story^37,38^) that can be compared between subjects. However, isolated choices are rarely meaningful units by themselves, as the choices that precede and follow them can change their interpretation. CBAS does not just evaluate the occurrence of individual choices, but, rather, the occurrence of all sequences up to a certain, user-defined, length. In general, the longer the sequences, the more data and computational resources will be necessary, but the richer the aspects of behavior that can be encompassed.

For the fly spontaneous alternation behavior, we applied CBAS to all sequences of choices of turns up to a length of 10. For each sequence of choices, all instances of that sequence up to a criterion of 250 choices are found in the dataset, and the rate of usage of that sequence in the two populations is used as the point of comparison at the basis of CBAS (Fig 1b shows one particular example sequence).

Correcting for the many comparisons inherent in evaluating the occurrence of such sequences is complicated by the fact that their incidence can be highly correlated. Standard methods to correct statistically for multiple comparisons (e.g. Bonferroni, Holm, Benjamini-Hochberg, Benjamini-Yekutieli) are underpowered for correlated data. Instead, we use an approach based on resampling to correct for the multiple comparisons—called the Romano-Wolf method. This approach retains power in the face of correlations^39^. In practice, we set the significance threshold according to median control of the false discovery proportion at 5%. This control is comparable to 5% control of the false discovery rate, which provides mean control of the false discovery proportion^40^.

Using the Romano-Wolf method for resampling-based multiple comparisons correction^39,40^, CBAS identifies many sequences of choices that differ significantly between the two strains of flies in the spontaneous alternation task (1,605/2,046 sequences; Fig 1c). We are also able to determine the sample size required to provide ∼80% power to detect a difference at this level of statistical certainty, using subsampling of the data (Fig S1).

### CBAS identifies interpretable differences for fly spontaneous alternation

CBAS identifies many sequences that significantly differ between the CA and w1118 fly lines. Upon inspection of all of the significantly different sequences, a clear, interpretable, contrast is apparent between those sequences that occur more in the CA line and those that occur more in the w1118 line (Fig 2a). The CA line utilizes sequences with extended numbers of the same turn in a row, whereas the w1118 line utilizes sequences with more frequent changes in the direction of turning. Therefore, CBAS not only identifies that there is a difference between the two fly lines, but it also provides information to support an interpretation as to the nature of the difference.

**Figure 2.**
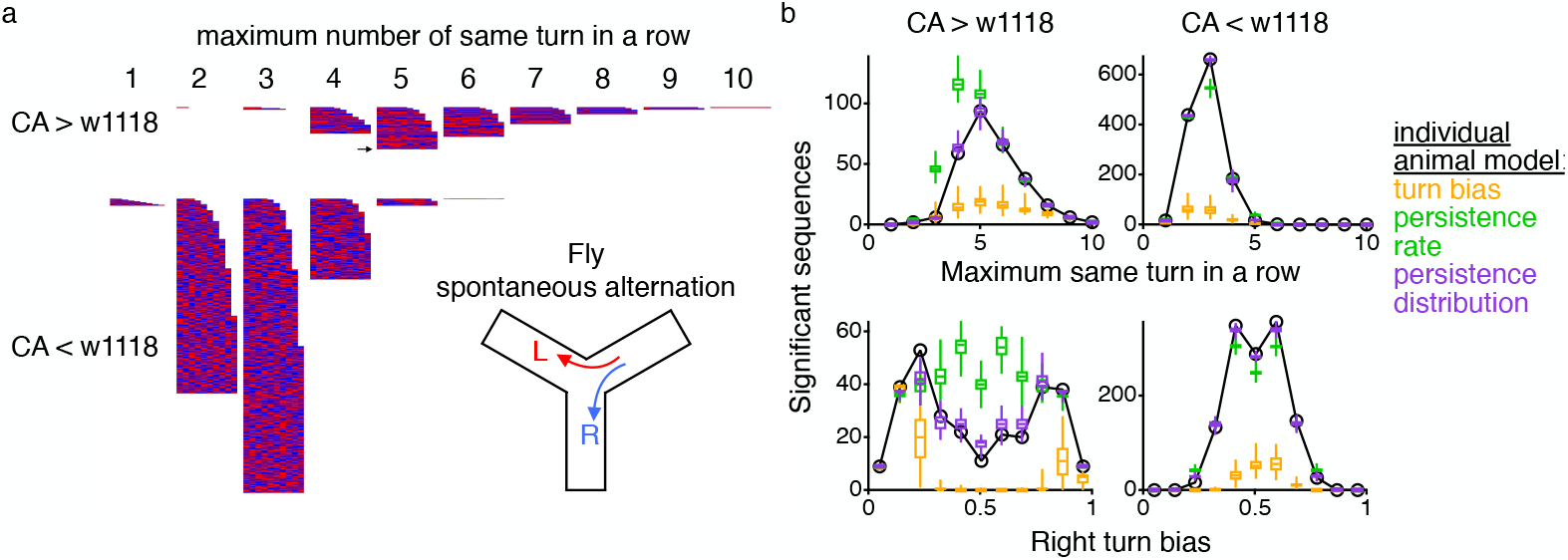
CBAS applied to fly spontaneous alternation produces interpretable differences between fly strains. CBAS was applied to flies tracked as they spontaneously made left and right turns on a y-maze. (**a**) All sequences up to length 10 identified as significantly different between the CA and w1118 strains are displayed with left turns indicated in red and rights turns indicated in blue. Each row displays a sequence. The top set of rows shows the sequences CA uses more than w1118 and the bottom set of rows are the sequences w1118 use more than CA. Sequences are ordered in columns based on persistence, i.e., the maximum number of the same turns in a row in the sequence. For example, the sequence displayed in figure 1b, RLRLRLLLLL, is in the column labeled 5 (indicated with an arrow) because there are 5 left turns in a row, and that is the maximum number of the same turns in a row in the sequence. (**b**) The number of significant sequences as defined by CBAS is displayed for the data (black line and circles) and three different models (colors and box and whisker displays) that might characterize the turns of the individual flies. The turn bias model (orange) randomly generates each turn with a bias based on the individual fly’s overall right turn bias over the first 250 turns. The persistence rate model (green) randomly generates each turn with a bias based on the individual fly’s overall rate of continuing in the same direction. The persistence distribution model (purple) randomly generates a sequence of the same direction of turns based on the individual fly’s probability distribution of continuing for any number of turns in a row. Top plots organize the sequences based on the maximum number of the same turns in row in the sequence (as in panel **a**). Bottom plots bin the same set of sequences based upon the fraction of right turns that occur in the sequence (30% for the example sequence in figure 1b). The left column quantifies the sequences that CA use more than w1118 and the right column the sequences w1118 use more than CA (or the equivalent model versions of each line). For the models, CBAS was run on 100 repeats of all three models, tailored to the choices of each individual fly. The center line of the box displays the median of the model output, the top and bottom lines of the box show the 75^th^ and 25^th^ quartiles, respectively, and the end of the whiskers show the full range of the model outputs.

The paper from which this open-source data comes from highlights the handedness of the individual flies, whereby individual flies show a propensity towards either right or left turns^31^. Such an observation creates a straightforward testable hypothesis about the nature of the difference between the two lines, i.e., that the lines differ in the extremes of their handedness bias, with individual CA flies hypothesized to being more extreme in their left/right turn bias than the w1118 lines. This hypothesis can be formalized by randomly generating virtual twin subjects by sampling turns according to the left/right turn biases of each individual fly. CBAS can compare this turn bias model of the lines, through generating an expectation for what its output would look like if the lines solely differed based on the left/right turn bias (Fig 2b). The CBAS results from the data are inconsistent with this hypothesis (judged by the numbers of sequences that are significantly different in two transparent metrics).

Instead of modeling the choices of the individual flies as being generated by a left/right bias on each turn, we could instead generate the turns based on the bias of the individual flies to persist in their direction of turning (i.e., turning in the same direction as their previous turn). If we generate virtual twin subjects by sampling turn directions according to these one-step persistence biases of each individual fly—persistence rate—the resulting CBAS output is closer to the data than the left/right turn bias but still is inconsistent overall (Fig 2b). However, if the choices of virtual twin flies are generated by using the distributions of persistence for each of the number of turns in a row from each individual fly—persistence distribution—then the CBAS of that model almost perfectly matches the CBAS output of the data according to our metrics (Fig 2b).

### Graceful decay of CBAS output with decreasing group size

We exploited the very large numbers of flies in the spontaneous alternation dataset to understand how the output of CBAS behaves with different sample sizes. This determines our ability to make conclusions and derive interpretations from fewer subjects. We first evaluate the number of significant sequences identified from smaller group sizes as a fraction of the total number of significant sequences identified with the full dataset (Fig 3a). As expected, with smaller group sizes, we find fewer overall significantly different sequences; however, across the range of group sizes evaluated, the median fraction of sequences was larger than the fraction of the population being used in the smaller group CBAS (Fig 3a). This indicates that at smaller group sizes, the proportion of sequences identified by CBAS for these subjects grows faster than the proportion of subjects in the CBAS. This provides a rapid increase in the amount of information provided by CBAS as the sample size increases.

**Figure 3.**
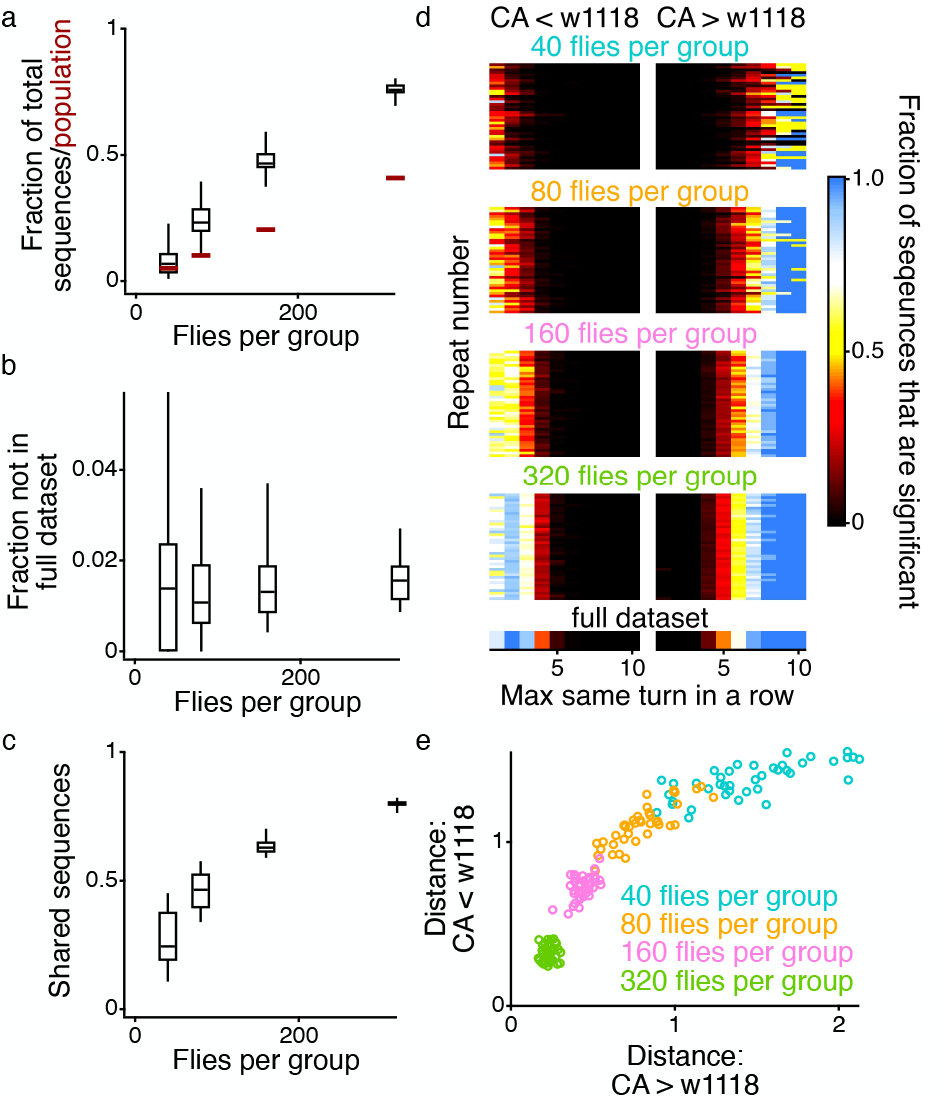
Scaling of CBAS with sample size. For all panels in this figure, 40 repeats of smaller sample sizes were generated from the fly data by selecting at random subjects within the two groups (without replacement). (**a**) Each CBAS run on the smaller sample size identified some number of significant sequences. Those sequences were compared to the sequences identified in the CBAS on the full dataset (with 759 CA and 807 w1118 flies), and the graph shows the ratios of the numbers of sequences identified in each instance of the smaller sample size that were also found in the full dataset to the total number of sequences identified in the full dataset. As a comparison, the ratio of the number of flies in the smaller CBAS to the total number of flies in the dataset is plotted in the maroon horizontal lines. (**b**) The ratio of the number of significant sequences identified in each instance of the smaller sample size that were not also identified in the full dataset to the total number of significant sequences identified in that instance of the smaller sample size. (**c**) In creating the smaller samples sizes, 20 paired sets of animals were generated such that the two groups of flies within a paired set had no overlapping individuals. Plotted are the number of sequences that were found to be significant in both groups of flies within a paired set of nonoverlapping flies over the average number of significant sequences found between the two paired set of flies. (**d**) For each instance of each sample size, all of the sequences were categorized based on the maximum number of turns in the same direction and the fraction of significant sequences within those categories are plotted to allow for an easier comparison across groups with very different numbers of sequences, i.e., there are only 2 sequences that have a maximum of 10 of the same turns in a row, whereas there are over 600 with a maximum of 3 of the same turns in a row. The bottom row of this plot is from the full dataset. (**e**) The Euclidean distance between each row from panel **d** and the full dataset row is plotted. Colors correspond to the sample sizes as shown in panel **d**. For panels **a** – **c** the center line of the box displays the median of the resamples, the top and bottom lines of the box show the 75^th^ and 25^th^ quartiles, respectively, and the end of the whiskers show the full range of the resampled outputs.

We next evaluate the fraction of the sequences identified by CBAS for the smaller sized groups that are not identified in the CBAS on the full dataset (which we might interpret as false positives). Since we control the false discovery proportion and not the familywise error rate, we do not expect this value to be zero; however, it was consistently small across all the group sizes evaluated (Fig 3b). The medians across the different repeats of the same sample sizes across the different sample sizes are all less than 2%.

Finally, we evaluated the similarity of the sequences identified by CBAS with nonoverlapping sets of subjects with the same group size (Fig 3c). At smaller groups sizes (40 per group), this overlap could be quite low (median 24.4%), even though there was sufficient evidence to discriminate the strains (Fig. S1). This means that the specific sequences identified by CBAS can be quite variable from one experiment to the next, especially at smaller group sizes. Given that, we sought to understand if the structure of the sequences comported with the conclusion from the full dataset—that the CA line uses more of the same turns in a row and the w1118 line more frequent changes in direction of turn. Even though the overlap between CBAS on different groups might not be that high (Fig. 3c), the sequences identified still follow the same structure as the full dataset in regard to number of the same turns in a row (seen across the repeats for each group size in Fig 3d). As the group size increases, the output of CBAS becomes more consistent, and more similar to the results from the full dataset (Fig 3e), licensing more generalized conclusions.

### CBAS can also correlate sequences of choices with a covariate of interest

The second task to which we apply CBAS is the two-step task, developed to test the interplay between model-based and model-free (reinforcement) learning algorithms^41^ (Fig 1a; middle). In this task, subjects choose between pairs of images at two different stages. The image choice at the first stage stochastically governs the pair of images from which the subject can choose at the second stage, and which enjoy drifting individual probabilities of delivering reward when picked. Model-dependent analyses of variants of this task (which make inferences based on the stochasticity in the transition between first and second stages) have led to conclusions about the algorithms used in different brains regions^41,42^ and differences that underlie psychiatric symptoms^43^, notably compulsivity. However, the interpretations are not without controversy^44^, and the complexity of behavior makes it difficult to evaluate ways in which the model might be missing features in the data.

Data for the two-step task come from a large population of human subjects in the open-source dataset from Gillan et al.^43^. We applied CBAS to all sequences of up to 4 choices long over the course of the 200 trials—400 total choices—of the task. Following Akam et al.^42^ we collapse image 3 and 4 into set A and image 5 and 6 into set B because the specific images within the set are not relevant for the critical decision at the first stage. Choices that are rewarded are treated as distinct from those that are not rewarded, as those choices can lead to different behaviors. That means that the sequence 2**A**1 is different from 2a1, where the upper-case emboldened letter indicates that a reward was received.

In building the two-step task dataset, the authors sought to examine how behavior varied along continuous trait dimensions. They did this by performing factor analysis on the answers to a series of psychiatric symptom questionnaires and showing that the way subjects performed that task was correlated with one of the factors (factor 2) that was associated with compulsive behavior and intrusive thoughts (CBIT)^43^. Therefore, we extended the CBAS so that it could also evaluate such correlations. We applied CBAS to the correlation between the sequence usage for each subject and their CBIT score (Fig 1b), finding 31/408 sequences to be significant (Fig 1c).

### CBAS provides evidence for testing model-dependent analyses of the two-step task

To understand the output of CBAS for the two-step task, we review the expectation associated with the hypothesis-driven analysis of this experiment. The two-step task was designed to evaluate the interplay between model-based and model-free decision making^41^ based on the stochastically rare transitions (from image 1 to set B; and image 2 to set A). Model-based decision making develops an understanding of the structure of the world (which is the model) and makes choices based on that understanding. Model-free decision making makes choices based on reinforced past successful actions, without developing an understanding of the structure of the world. A way to see the difference between these two decision-making schemes is when a learner chooses image 2 at the first step, makes an unlikely stochastic transition to set A, and gains a reward at this second step (‘2**A**’ in our language). A regular model-free learner will reinforce the choice of image 2 (along with the image chosen at A) and will therefore be more likely to choose image 2 on the next trial. By contrast, a model-based learner will be more likely to choose image 1 on the next opportunity, based on the observation that this will make the transition to set A more likely (because of the common/rare transition structure in the environment)^41^.

We evaluated all of the sequences that CBAS found as either significantly positively or negatively correlated with CBIT (Fig 4a). CBAS identified a positive correlation with many sequences involving being rewarded at A and then selecting image 2 or being rewarded at B and then selecting image 1. In particular, the structures 1**A**2 or 2**B**1 occur commonly in the positively correlated sequences. These behavioral motifs are inconsistent with either model-free or model-based learning, as the subjects get rewarded after choosing from the common side (A from 1 or B from 2) but then select the image that will rarely bring them back to the previously rewarded side (2 from A or 1 from B). The model-based decision maker would return to 1 as that is the side that would commonly go to set A where there was the reward. The model-free decision maker would also return to 1 as that was the side that was reinforced as being connected to the recent reward at A.

**Figure 4.**
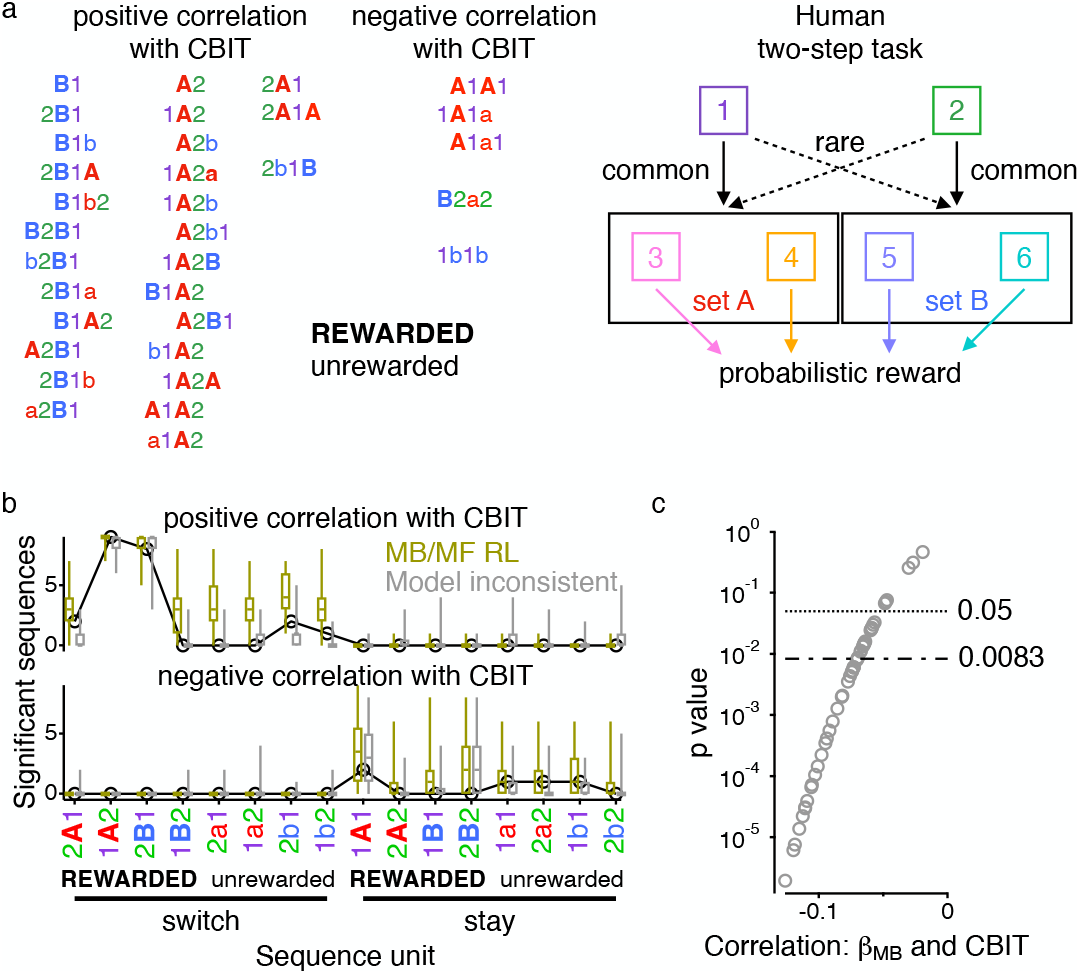
CBAS helps interpret the correlation between CBIT and the two step task. CBAS was applied to humans performing the two-step task with sequences of choices up to length 4. (**a**) All sequences whose usage was positively (left) and negatively (right) correlated with the individual subject CBIT factor score are displayed. Colors correspond to the images and image sets chosen as indicated by the diagram of the task (right). Upper case bold letters indicate that a reward was received. The majority of the sequences positively correlated with CBIT include **B**1 and/or **A**2, and so the first two columns are arranged to center these motifs. For illustrative purposes, the two sequences **B**1**A**2 and **A**2**B**1 are duplicated and listed in both columns. (**b**) The number of significant sequences as defined by CBAS is displayed for the data (black line and circles) and two different models of the task (colors and box and whisker displays). Sequences are organized based upon their containing the subunit indicated on the x-axis. MB/MF RL model (yellow) is the model-based/model-free RL model from Gillan et al.^43^. Each subject was fit to the model, and their parameters were used to generate choices for CBAS. Model inconsistent choices (grey) were generated by using the fit parameters from the MB/MF RL model for a single subject, but whenever the sequence 1A or 2B was chosen, the subsequent choice was probabilistically determined based on the “model inconsistent” rate of the individual subject. The “model inconsistent” rate for a subject was their rate of choosing a 2 following 1A or a 1 following 2B. For each CBAS repeat of the model inconsistent choices a different subject’s fit parameters were used. For the models, CBAS was run on 100 repeats. The center line of the box displays the median of the model repeats, the top and bottom lines of the box show the 75^th^ and 25^th^ quartiles, respectively, and the end of the whiskers show the full range of the odel repeats. (**c**) Data generated from the sequences of choices matched to the model inconsistent rate of each subject (as described above and shown in grey in panel **b**) was fit to the MB/MF RL model and the p-value of the correlation of the model based parameter of the model fit with CBIT is displayed as function of the correlation value. Dotted horizontal lines show p = 0.05, and p = 0.0083. The latter number is the Bonferroni corrected p-value to control for the 6 parameters of the model. Bonferroni correction is used instead of Romano-Wolf, as that would be the most typical, if not conservative, correction used. A total of 50 model repeats was fit, 86% and 56% had p-values <0.05 and 0.0083, respectively.

We next used CBAS to test whether the combined model-based/model free reinforcement learning (MB/MF RL) model used by Gillan et al.^43^ which has parameters describing model-based and model-free decision making and the propensity to employ each, correctly captures the data. To do this, we generated virtual twin choices for each subject based on fits of each subject to this model and ran CBAS on the model output (Fig 4b). This led to a positive correlation between CBIT and the model inconsistent sequences with the choices 1**A**2 and 2**B**1, like the human data. However, there was the important difference in that CBAS identified a positive correlation with the unrewarded versions of those units (1a2 and 2b1) that was not present in the human data.

CBAS therefore identified an important failing in the MB/MF RL model. We can explain this by noting that this model has a parameter for “stickiness”—the propensity to continue to choose the same of the first images (1 or 2) in a row. Decreased stickiness would also comparatively favor sequences that change the first image, including 1**A**2, 1a2, 2**B**1 and 2b1. Indeed, in our fits, along with the negative correlation between the propensity to employ model-based choices and CBIT (*ρ* = −0.08; *p* = 1.7*x* 10^-3^) reported by Gillan et al.^43^, there was also a negative correlation between the stickiness parameter and CBIT (*ρ* = −0.11; *p* = 1.6*x* 10^-5^).

As an alternative, we generated behavior based on just model-inconsistent choices. That is, we created a virtual twin of each subject s from the MB/MF RL model by hybridizing the rate with which that subject performed model-inconsistent choices with choices that were otherwise generated according to the fit parameters for a single, reference subject s*. The reference subject s*, was randomly chosen for each of the 50 repeats. That is, we used the MB/MF RL model based on s* to generate data, but every time the model produced 1**A** or 2**B** we probabilistically selected the next choice according to the overall model-inconsistent rate of subject s (but then continued to generate choices as if nothing else had changed). This ensures that the majority of choices came from a subject with a single CBIT value, and only the choices following 1**A** or 2**B** were manipulated (19% on average). The CBAS results from this model-variant were much closer to the CBAS output of the data in the metric reported (shown in the ‘Model inconsistent’ results in Fig. 4b). Furthermore, when we fit the output of the model-inconsistent variant of the model to the MB/MF RL model, we found that a positive correlation between model-inconsistent choices and CBIT alone can appear as a negative correlation between the model-based parameter of the model and CBIT (Fig 4c).

### CBAS identifies a hippocampal dependent strategy for spatial alternation learning

The third, and final task we evaluated with CBAS is spatial alternation behavior in rats. For this task, to get reward, the rats must alternate between pairs of arms of a track whilst visiting a different arm of the track in between (Fig 1a; right). Spatial alternation is a common behavioral paradigm for phenotyping and neurophysiology, and the discretization of the behavior into the arms chosen by the animals forms the basis of many of the conclusions from these studies^45–49^. However, our recent work calls into question the assumptions and hypotheses that motivate the standard analysis for spatial alternation behavior^50,51^. Such limitations could even be general across other tasks used to evaluate memory^52^. Furthermore, even though we have developed many variants of reinforcement learning agents to fit individual behavior, these agents learned in ways that were very evidently different from the animals^51^, limiting their use for phenotyping. Here, we use CBAS to evaluate the role of the hippocampus in this task by comparing rats with lesions to their hippocampus to rats that underwent control surgery.

For the spatial alternation task, we applied CBAS to all sequences of arm choices up to 6 choices long over the first 800 arm choices of the rats. As with the two-step task, rewarded choices are treated differently from unrewarded choices. As with the fly spontaneous alternation dataset, CBAS compared the rate of usage of each sequence between the populations of hippocampally-lesioned and control rats (Fig 1b; right). 409/24,342 sequences differed significantly (Fig 1c).

CBAS identified many sequences discriminating the fly strains. However, the dimensionality of the task is quite low, so it is fairly straightforward to structure all of the sequences to develop an interpretation (Fig 2). By contrast, CBAS did not identify that many sequences in the two-step task that significantly correlated with CBIT, so the extra complexity and dimensionality of this task did not prevent us from directly evaluating all the sequences to develop an interpretation (Fig 4). For the case of rat spatial alternation, CBAS identified a good number of sequences, and the task is complex and high dimensional, making it a challenge to directly develop an interpretation of the output of CBAS.

Therefore, we developed an analysis method that takes advantage of the structure of the sequences at the output of CBAS to help inform an interpretation of the spatial alternation behavior. CBAS evaluates all sequences up to a certain length, and so sequences with shared and overlapping structure might all be significant. We therefore defined the notion of a “complete” target sequence (Fig S2) as one with the property that all sufficiently prevalent longer sequences that contain it are found by CBAS to differ significantly in the same way as the target (e.g., all are found to be greater in the control than the lesion rats). Effectively, a “complete” sequence has the property that no matter what happens before or after that sequence, the series of choices remain significantly different between the groups. Such sequences are a powerful way to simplify the complex behavior.

We find “complete” sequences that the control rats do more than the lesion rats and that the control rats do less than the lesion rats (Fig 5a). Inspection of these sequences provides two axes to compare all of the significant sequences. The “complete” sequences that the control rats use more than the lesion rats show systematic exploration of the arms by largely continuing in the same direction and transitioning mostly to neighboring arms. These features are much less the case in the “complete” sequences that the control rats do less than the lesion rats. We used these two axes to compare all the significant sequences (Fig 5b). Consistent with the finding for “complete” sequences, sequences that the control rats use more than the lesion rats tend towards higher directional inertia—the propensity to continue in the same direction— with smaller overall transition distance.

**Figure 5.**
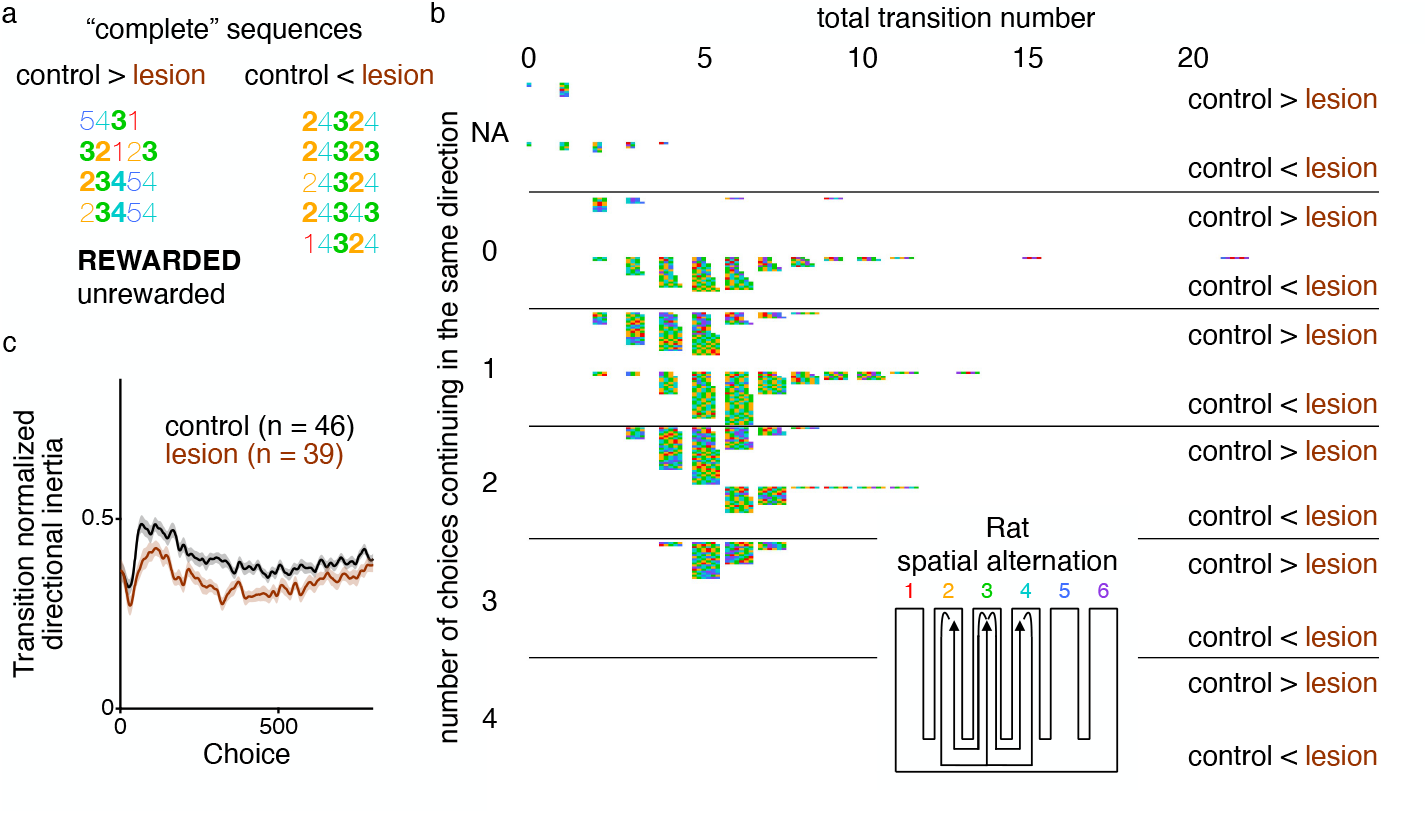
CBAS identifies hippocampal based learning strategy for spatial alternation task in rats. CBAS was applied to rats performing the spatial alternation task with sequences of choices up to length 6. (**a**) “Complete” sequences that the control rats use more than hippocampal lesion (left) and that the lesion rats use more than control (right). “Complete” sequences are ones that that always significantly different between the groups (in the same direction) no matter what choices occur before or after up to the maximum length of 6 (see Fig. S2). Emboldened numbers indicate that choice was rewarded. (**b**) The “complete” sequences differ between the groups in two broad ways: the consistency of the direction of travel (greater for controls); and the total absolute distance covered (so 21 would contribute a distance of 1 and 14 would contribute a distance of 3). This plot organizes all the sequences (i.e., not just the “complete” sequences) according to these two features (the former defining the rows, with NA for sequences with fewer than three choices for which this cannot be calculated; the latter the columns). For example, sequence **32**12**3**, is located in the row segment labeled 2 as two of the choices in the sequence continue in the same direction: **3**-**2**-1 (the 2-1 is in the same direction as 3-2) and 1-2-**3** (the 2-3 is in the same direction as 1-2), and is located in the column segment labeled 4, as all of the choices are 1 arm away from the previous choice. Sequences that are 1 choice long have a total transition number of 0. Each sequence in the plot is displayed along the tiny rows ordered by length, and, for completeness, the choices are colored based on the color of the arm numbers in the task diagram. (**c**) Given that the sequences that occur more in the control than the lesion rats tend to have more choices that continue in the same direction and smaller total transition numbers, we captured these feature of the data using the metric of transition normalized directional inertia, which measures the tendency to continue in the same direction divided by the distance of the transition. Transition normalized directional inertia was smoothed and averaged (±sem) across all rats in the control (black) and lesion (brown) groups and shown for the first 800 choices of the spatial alternation task. Control rats show significantly larger transition normalized directional inertia summed across the 800 choices of the spatial alternation task, p = 4.0 x 10^-4^ (permutation test).

The hypothesis generated by the “complete” sequences and verified across all of the significant sequences, motivates a relatively straightforward metric by which we can evaluate all of the choices of the rats for comparison between the two groups. We quantified the transition normalized directional inertia in the control and lesion rats. This metric measures, for each choice, whether that choice continues in the same direction as the preceding choice and normalizes it by the number of arms transitioned. For example, choosing arm 3 after coming from 2 and 1 (sequence 1-2-3) has a transition normalized directional inertia of 1, choosing arm 4 after coming from 2 and 1 (sequences 1-2-4) has a transition normalized directional inertia of 0.5, and choosing arm 1 after coming from 2 and 1 (sequence 1-2-1) has a transition normalized directional inertia of 0. Lesion and control rats show a clear and significant difference in their transition normalized directional inertia (Fig 5c).

Although “complete” sequences were not used to interpret the fly and human CBAS outputs, evaluating them would lead to identical conclusions (Fig S3).

### CBAS generated hypothesis replicates for spatial alternation

The fact that the transition normalized directional inertia is significantly different between the control and lesion rats, is partly a circular claim. We motivated the metric based on the sequences identified by CBAS as differing between the groups. It did not have to be the case that the metric would differ across all the choices of the animals (as opposed to just the significant sequences), but it should not be overly surprising that it does. Therefore, we sought to replicate our understanding in a separate dataset of hippocampal lesion and control rats.

Even with a smaller sample size, CBAS identifies differences due to the hippocampal lesioning (Fig 6a), consistent with the power calculation on the initial dataset (Fig S1e&f). The sequences identified by CBAS separate along the two axes identified by the “complete” sequences from the initial dataset (Fig 6b). This comportment of the structure of the sequences with a smaller sample size is consistent with what was found with subsampling the fly data (Fig 3d). Consistent with the hypothesis from the initial dataset, the groups significantly differ in their transition normalized directional inertia (Fig 6c). Indeed, we even replicated specific sequences across the two datasets (Fig 6d). Interestingly, a much larger fraction of the sequences that the control rats use more than the lesion rats replicate compared to the sequences the control rats use less than the lesion rats. This is consistent with the overall hypothesis that the hippocampus enables a learning strategy that systematically engages with the arms of the track through moving in the same direction and transitioning to nearby arms. Without a hippocampus that systematic behavior goes away, exposing less systematic choices, that inevitably will be less likely to replicate between samples.

**Figure 6.**
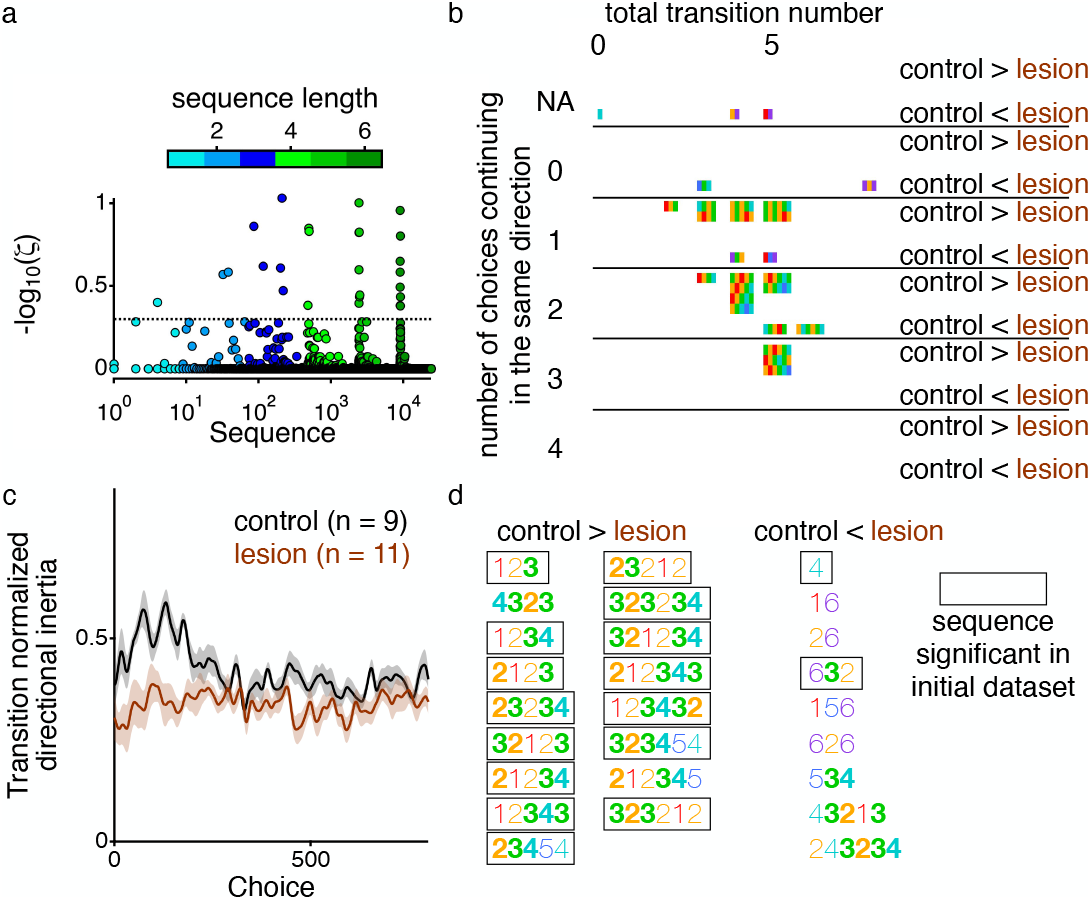
Replication of the CBAS generated hypothesis in the rat spatial alternation task. CBAS was applied to a separate set of control and hippocampal lesion rats performing the spatial alternation task. (**a**) CBAS Manhattan plot displaying the p-value for each sequence compared between the two groups. Plot structured as described in Fig. 1c. (**b**) Significant sequences structured by the two metrics, number of choices in the same direction and total transition number, as described in Fig. 5b. (**c**) Transition normalized directional inertia as the two groups of rats in the replication dataset learn the spatial alternation task. Control rats show significantly larger transition normalized direction inertia summed across the 800 choices of the spatial alternation task, p = 2.2 x 10^-3^ (permutation test). (**d**) All of the significant sequences that CBAS identifies as occurring more in the control than lesion rats (two left columns) and as occurring less in the control than lesion rats (rightmost column). 65.4% (88.2% of for control > lesion, 22.2% of for control < lesion) of the sequences CBAS identifies as significant in this dataset were also significant in the initial dataset (indicated by boxes).

## DISCUSSSION

We have presented a general data-driven behavioral analysis method, CBAS, for identifying interpretable behavioral components that is grounded in sequences of choices made by subjects. There has been significant progress, in recent years, in tracking and analyzing, short timescale actions of subjects^7–16^; however, we lack methods for generally analyzing and interpreting sequences of these actions or choices and decisions of subjects that unfold over a longer period than moment to moment movements. CBAS provides just such a method and is applicable across a wide array of species and different behavioral paradigms (Fig 1). It can be used to test models and hypotheses (Fig 2&4), and, for instance, using generalizable simple principles about the relationship between sequences, it can generate hypotheses in complex behaviors where reliable computational understanding has yet to emerge (Fig 5&6). Through taking advantage of large-scale data collection and rigorous statistical methods, CBAS has the potential to transform our use of behavior in a comparable way to how GWAS/WES/WGS changed the paradigm for genomic studies.

Somewhat surprisingly, in all tasks, CBAS provides information and generates interpretations that are either inconsistent with, or unexpected from, the prior understanding of these data and tasks. For the flies, as mentioned above, the dataset came from a paper characterizing the existence of handedness in these flies, as manifested by extremes in individual preferences for left or right turns^31^. The output of CBAS on these data allowed for the testing of generative models to explain that handedness. The most straightforward model that one could postulate based on the hypothesis of handedness is not consistent with the data (Fig 2b); whereas a model drawing upon the distribution of individual flies to persist in the same direction for a range of the same turn in a row was almost indistinguishable from the data according to our metrics. Therefore, CBAS generates testable hypotheses as to underlying computational structure that these flies draw upon as they perform this behavior.

For the two-step task, the CBAS results provide an alternative interpretation for the relationship between the task and individual CBIT levels. Previously, these data were interpreted to mean that individuals with more CBIT are less likely to draw upon model-based planning^43^. The CBAS results indicate that the relationship with CBIT rests more on the increase of model-inconsistent choices with CBIT (Fig 4a&b), and that that increase alone could appear as a negative correlation with model-based planning, even if there was in reality not one (Fig 4c). What an increase in model-inconsistent choices means in the context CBIT is something that should be explored in the future.

The result that the hippocampus is involved in spatial alternation behavior is not a new one. There have been prior reports of changes to spatial alternation behavior in the context of hippocampal lesioning^45^ and manipulation^48,53^. However, all of the prior work occurred from the prevailing perspective of spatial alternation exclusively reflecting memory allocation, something we have recently shown to be inconsistent with the way in which the rats learn^50,51^. Guided by the results of CBAS we find that the hippocampus is responsible for implementing a specific strategy (or set of strategies) whereby the animals systematically move across the track and transition between closer arms as they learn. This strategy is far more consistent with the hippocampus implementing a structuring of the task space as described in recent models of hippocampal function^54,55^, than with the typical memory function assumed in the prior studies of spatial alternation behavior.

These three tasks were just the first ones to which we applied CBAS; they were not selected from a larger set of tasks and used for illustrative purposes. We strongly anticipate that there will be comparable benefits to applying CBAS and other data-driven methods to a broad array of datasets and behavioral tasks to provide a stronger foundation for our understanding of behavior. The ability of CBAS to identify differences or correlations in a dataset is directly built into its function. However, the ability to interpret the differences or correlations found remains nontrivial. We have provided examples of ways to extract interpretable information from the sequences output by CBAS and have developed a general analysis (“complete” sequences) to assist in that process. However, there remains room for future work to assist in going from the identification of a difference or correlation by CBAS to understanding and interpreting what those differences and correlations mean. Of course this is in no way a unique problem to CBAS. Genomics methods (GWAS/WES/WGS) similarly require secondary and follow up analyses.

### Considerations for the application of CBAS to datasets

To apply CBAS to a dataset, there are some decisions that need to be made. At a fundamental level, CBAS requires a discretized description of the behavior. This was motivated by the common use of choices to describe behavior (Fig 1a). Furthermore, there is evidence that, at a rather fundamental level, behavior occurs through discrete choices^12,56^. CBAS is not currently capable of working on continuous data. The discretization informs the language that is used to make up the sequences that form the basis of the statistical evaluation within CBAS. In deciding on the language there is a range of biases that can be imposed with different decisions. For example, with the two-step task, we combined the second image choice into sets A or B. This prevents CBAS from differentiating a difference in choices at that level. Based on the literature we deemed that bias to be justified, but it is a bias, nonetheless.

A next decision to make is the length of sequence up to which CBAS will evaluate. The longer the sequences, the more information CBAS can provide, but the more data and computational resources that will be necessary. If the language is large, i.e. there are many things that can happen in the behavior at any point in time, and the transition matrix between one choice and another is not sparse, then computational resources might not be feasible to calculate the statistics underlying CBAS, especially if longer sequences are evaluated. Additionally, it is necessary to specify a criterion for the total number of choices per animal or session that are decomposed to form the sequences that are compared. Although it is possible just to include all the data, this might introduce bias if the groups have different total numbers of choices overall. Conversely, one might try and study aspects of learning or the acquisition of competence by applying CBAS to restricted portions of the behavior. It would also be possible to use a discrete language that characterizes aspects of behavior that unfold over longer timescales (for instance, defining a discrete language using unsupervised clustering over windows of choices), or indeed to employ a multi-scale description.

Finally, CBAS either needs to compare groups or correlate with a covariate of interest. Currently CBAS is not able to just evaluate a single group of animals by itself in an unsupervised manner without a covariate of interest. Future work will focus on relaxing this limitation.

In sum, we offer CBAS as a new data-driven analysis technique that extracts statistically rigorous differences in behavioral sequences between or among groups. We show that such sequence distinctions can be revealing about the underlying information processing functions that vary and can circumvent biases that plague model-dependent analysis methods. We illustrated the properties and promise of CBAS in three different datasets in three different species.

## Methods

### Romano-Wolf resampling based multiple comparisons correction

We follow the terminology and description laid out in Clarke et al.^57^ to describe the Romano-Wolf multiple comparison correction. First, we describe the way the method corrects for the familywise error rate (FWER) and then explain how the procedure is extended to provide median control of the false discovery proportion. FWER control at a level of *α* means that across all comparisons there is a *α* percent chance of having at least one false positive rejection of a null hypothesis. The Romano-Wolf procedure provides FWER control through resampling the data^39^. We describe the case of testing a total of *S* hypotheses.

It is not generally known if the Romano-Wolf procedure controls for type III, or directional, errors. Type III errors are errors in the sign, or direction, of the conclusion. For example, if a statistical test provided information to reject the null hypothesis *θ*_1_ = *θ*_2_, and you then concluded that *θ*_1_ > *θ*_2_, when in fact *θ*_1_ < *θ*_2_. Therefore, instead of running a single two-tailed test for each sequence, we run two one-tailed tests for each sequence. As a consequence, the total number of hypotheses tested, *S*, is twice the total number of sequences being compared. For CBAS those hypotheses take one of two forms: i) the rate of each sequence (*r*_*s*_) is the same between two groups Δ*r*_*s*_ = 0, or ii) that there is no correlation (*ρ*_*s*_) between each sequence and a covariate of interest, *ρ*_*s*_ = 0. For the one-tailed versions of each hypothesis, we ask if 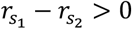 and 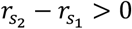 for the comparison CBAS, where 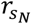 is the rate of sequence *s* for group *N*, or if *ρ*_*s*_ > 0 and *ρ*_*s*_ < 0 for the correlational CBAS.

The first step in the procedure is to create a studentized test statistic for each hypothesis. The studentization is different based on whether the CBAS is comparing two groups or calculating a correlation. In the case where two groups are being compared:

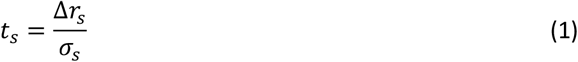

where *σ*_*s*_ is the standard error of Δ*r*_*s*_, which we calculate by combining the standard error of the mean of the rate for each group using error propagation, i.e. 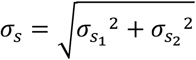 where 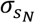 is the standard error of the occurrence rate of sequence *s* for group *N*

In the case where the correlation is being calculated the studentized test statistic is^58^:

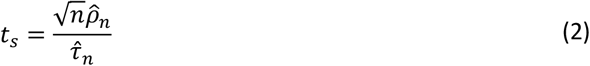

where:

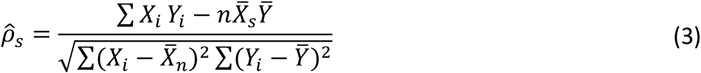

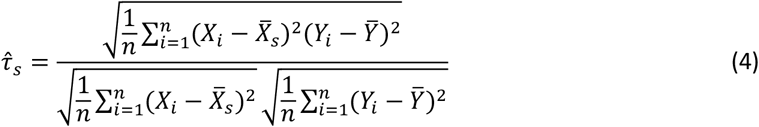

For eq. 2, 3, and 4, *n* is the number of subjects for which the correlation is being calculated, *X*_*i*_ and *Y*_*i*_ are the values of the metrics being correlated (in our case, the sequence count and CBIT score for each individual respectively) 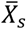 is the mean sequence count for the specific sequence being considered, and 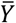 is the mean of the covariate of interest (CBIT score, which is the same for any sequence).

When two groups are being compared, we resample from the entire population with replacement (separately for each group) and build up a null distribution by bootstrapping *M* times. The test statistic from the *m*^*th*^ bootstrap sample for *m* =1,…*M is:*

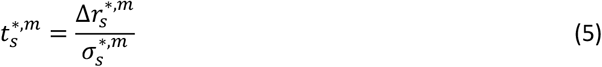

Where, 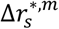 is the difference in the rate of each sequence whilst resampling, with replacement, from the entire population, ignoring the group labels 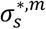 is the accompanying bootstrap standard error of the difference in means. The resampled group sizes are the same as the two groups of interest.

In the case where the correlation is being calculated, the test statistic based on the *m* ^*th*^ bootstrap sample is:

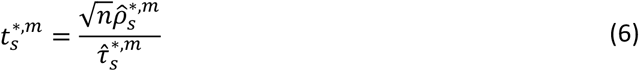

where 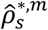 is the correlations of each sequence whilst permuting the relationship between the metrics being calculated across the entire population 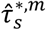 is the accompanying normalization for the permuted relationship. The permuted group size is the same as the original.

For the main CBAS calculations, we used a value of *M* = 10,000. When we run many repeats of different models or for power calculations, we use a value of *M* = 1,000. Importantly, for each individual resampling, *m*, the same resampled set is used for all sequences.

The test statistics, and their accompanying estimators are ordered from largest to smallest values. This creates an *M* × *S* matrix where each column contains all the estimators of the test statistics. The first column contains the estimators from the largest test statistic, the second column contains the estimators from the second largest, etc.

To define the distribution for which each test statistic is compared, which then determines the adjusted p-value, the following algorithm is used. The first sequence considered is the one with the maximum test statistic, *t*_*s*_. Its comparison distribution is defined as the maximum value within each row of the matrix of estimators of the test statistic:

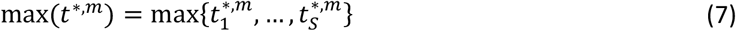

which provides a total of *M* values, *t* ^* *m*^(there is no longer an association with *s*, because these values can come from a resampling of any of the sequences). Using those *M* values, the adjusted p-value is calculated as follows:

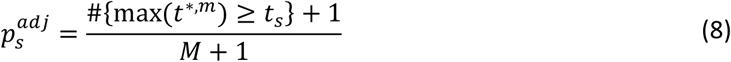

After calculating 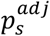 for the first sequence, the column with the test statistic estimators generated from the first sequence is removed from the matrix. This now leaves a matrix that is *M* × (*S* − 1). The above procedure is then used to calculate 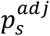 for the sequence with the second largest test statistic, and then its column of test statistic estimators is removed, etc until all 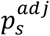 have been calculated. Following the algorithm described in Clarke et al.^57^, we enforce monotonicity of the p-values by resetting the p-value for each sequence:

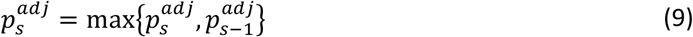

This is done prior to calculating the p-value for the next sequence.

To control the false discovery proportion (i.e., control the number of false positives divided by the total number of hypotheses rejected), the idea of k-FWER is introduced^40^. For control of the FWER, k is equal to 1, and that leads to an *α* percent chance that there is at least 1 false positive among all hypotheses rejected (for FWER control *α* = 0.05 is most commonly used). If k equals 2, then there is an percent chance that there are at least 2 false positives among all hypotheses rejected. Therefore, to get control of the false discovery proportion we need to find the k that provides the proportion of interest given the number of hypotheses rejected. So, if we want a false discovery proportion, *γ*, of 0.05, we need *k*∼0.05 × number of hypotheses rejected.

Romano and Wolf also derived an algorithm to do just that. The algorithm is as follows. Start with *k* = 1. Apply the k-FWER procedure, and note the total number of hypotheses rejected *N*. If 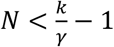 stop and you have identified k. Otherwise, increase k by 1, and repeat^40^. The way you determine k-FWER is in eq. 7, instead of taking the maximum value in each row, you take the k^th^ largest value. Finally, to get median control of the false discovery proportion *α* = 0.5. This means that 50% of the time you will get a value greater than *γ*, and 50% of the time you will get a value less than *γ*, leading to median control of the false discovery proportion^40^. This is a similar decision to what is done when calculating the false discovery rate (FDR) with Benjamini-Hochberg or Benjamini-Yekutieli, except the false discovery rate controls the mean of the false discovery proportion instead of the median.

Lastly to avoid confusion around what the statistical control is we label the adjusted p-value with the k-FWER control (i.e. median control of the false discovery proportion) as *ζ*instead of p, comparable to the use of q with FDR control.

### Power estimation for sample size calculation with CBAS

To estimate the statistical power of CBAS for a given sample size (Fig S1) we resampled the dataset without replacement and ran CBAS to determine the number of significant sequences. For each sample size we performed 20 repeats. We also ran a CBAS comparing each group to itself, with 20 repeats for each group; or correlating the sequence counts with a randomly generated set of CBIT scores drawn from the same distribution as the actual CBIT scores (Fig S1), with 40 repeats. Then for each sample size, the power is estimated by identifying the fraction of largest 20 number of significant sequences that came from the CBAS run on the resampled data (as opposed to the comparisons comparing groups to themselves or randomly generated CBIT).

### Power estimation for sequence usage for “complete” sequences

Populations the same size as the datasets were bootstrapped (sampled with replacement). 1,000 CBAS were run on different bootstrapped samples. The smallest sequence usage rate for which there was 80% power to detect a difference was defined as the smallest usage rate from the sequences that were significant on 80% of the CBAS runs.

### Spontaneous alternation in flies

#### Data

Data was downloaded from: https://lab.debivort.org/neuronal-control-of-locomotor-handedness/. The first 250 turns for each fly were used. Flies that did not do at least 250 turn were excluded from the analysis. 466/1,225 CA and 565/1372 w1118 flies did not reach the 250-turn criterion.

### Two-step task in humans

#### Data

Data was downloaded from: https://osf.io/usdgt/overview. All subjects in the dataset completed all 200 trials/400 choices. Therefore, all subjects were included in the analysis.

#### MB/MF RL model

The model was implemented following the description from Gillan et al.^43^. The model combines units for model-based and model-free control, as well as stickiness. For each trial, *t*, there are choices at two stages, *c*_1,*t*_ and *c*_2,*t*_. The choice at stage 1 leads stochastically to a stage 2 state, *s*_*t*_. Reward, *r*_*t*_, occurs following the second stage choice based on dynamic probabilities for each image at the second stage. *r*_*t*_ = 1 indicates reward and *r*_*t*_ = −1 indicates no reward. Stage 1 choices depend upon what is learned at stage 2; therefore, stage 2 learning is described first.

At stage 2, the model learns a value function, 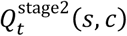 where *s* is state. The reward received at each trial leads to updating *Q* according to: 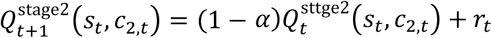 where *α* is the learning rate parameter. The probability to make a specific stage 2 choice is governed by a softmax: 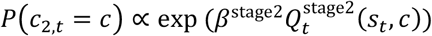 which is normalized over both options of c, and *β*^stage2^ is the inverse temperature parameter.

Stage 1 choices are determined by weighting the value predictions of the different components of the model about the stage 2 value of each stage 1 choice. Model-based values are dictated by the learned values of the stage 2 state, maximized over the two actions: 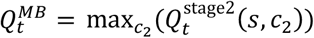 where *s* is the stage 2 state predominantly produced by stage 1 choice *c*. Model-free values are dictated by two learning rules: 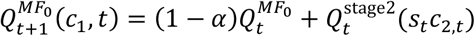 and 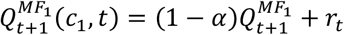 Stage 1 choice probabilities are also governed by a softmax: 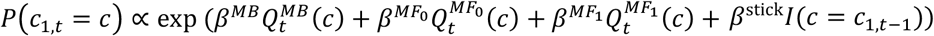 where the *β* values are the temperature parameters for each component, and *I*(*c* = *c*_1,*t*−1_) is 1 for the choice that repeats the one made on the previous trial and 0 for the other choice.

Following each trial, the value estimates in all *Q* for all unchosen actions and unvisited states are decayed multiplicatively by (1 − *α*).

Simulated annealing was used to fit the model, initializing with random parameters. At least 10 different fits were run for each set of parameters and the fit parameters leading to the largest log-likelihood was chosen. The data contained instances where a subject did not choose either of the images presented. Those choices were not included in the likelihood for the fitting. However, the lack of choice was coded as its own symbol and analyzed within CBAS for correlations with CBIT.

### Spatial alternation in rats

#### Animals

All experiments on rats were conducted in accordance with University of California San Francisco Institutional Animal Care and Use Committee and US National Institutes of Health guidelines. Rats were fed standard rat chow (LabDiet 5001). To motivate the rats to perform the task, reward was sweetened evaporated milk: 25 g of sugar per can (354 ml) of evaporated milk (Carnation). The rats were food restricted to up to 85% of their basal body weight.

#### Surgery

Anesthesia was induced using ketamine, xylazine, atropine, and isoflurane. The animals underwent surgery to either lesion the dorsal and intermediate hippocampus or as control for the lesioning. For the lesion rats, craniotomies were drilled bilaterally to allow for access to all 24 injection sites (Table 1) and the injection needle (33 gauge) was inserted through the dura. For each lesion site, the needle was lowered 0.1 mm below the target coordinates and then raised back to the target prior to injection. 120 or 110 nl of NMDA (20 mg/ml) was injected at each site at a rate of 0.1 µl/min, for males and females, respectively. The needle was left at the site for 2 minutes after finishing the injection. For the rats that underwent control surgeries, everything was the same as the lesion rats, except no NMDA was injected, and the needle did not dwell at the hypothetical injection site. All animals that underwent the lesion surgery received Diazepam 10 mg/kg intraperitoneally post-operatively.

**Table 1.** Stereotaxic coordinates of NMDA infusions to produce lesions within the hippocampal formation. Adapted from Kim *s* Frank^45^. The coordinates are given for a Long Evans rat skull which is leveled so that bregma and lambda lie in the same horizontal plane. AP: anteroposterior, ML: mediolateral, DV: dorsoventral. The AP and ML coordinates are measured from bregma. DV coordinates are measured from the dural surface above the brain.

| AP (mm) | ML (mm) | DV (mm) |
| --- | --- | --- |
| -2.8 | ±1.4 | -3.0 |
| -3.3 | ±2.4 | -3.0 |
| -4.1 | ±1.8 | -2.8 |
| -4.1 | ±3.4 | -2.8 |
| -4.8 | ±2.0 | -2.8 |
| -4.8 | ±4.2 | -3.1 |
| -5.5 | ±2.6 | -3.0 |
| -5.5 | ±3.6 | -2.9 |
| -5.5 | ±5.0 | -3.5 |
| -5.5 | ±5.0 | -5.5 |
| -6.2 | ±4.0 | -3.4 |
| -6.2 | ±5.4 | -4.4 |

Hippocampal lesioning was confirmed via micro-CT (Fig S4) collected at the Advanced Light Source at Lawrence Berkeley National Lab. Brains were processed and imaged, and scans were processed as described in Kastner et al.^59^. CT images were aligned using manual registration in Slicer-3D (http://www.slicer.org)^60,61^. Male and female brains were separately registered.

#### Spatial alternation behavior

The automated behavior system for spatial alternation behavior was previously described^51^. There are different symbols on each arm of the track serving as proximal cues, and there are distal cues distinguishing the different walls of the room. Pneumatic pistons (Clippard) open and close the doors. Python scripts, run through Trodes (Spike Gadgets), control the logic of the automated system. The reward wells contain an infrared beam adjacent to the reward spigot. The automated system uses the breakage of that infrared beam to progress through the logic of the behavior. In addition to the infrared beam and the spigot to deliver the reward, each reward well has an associated white light LED.

Each cohort of rats is divided into groups of four (or three) animals. The same groups were maintained throughout the duration of the experiment. Within a group, a given rat is always placed in the same rest box, and the four rats of a group serially perform the behavior. The rats have multiple sessions on the track each day. Prior to beginning the spatial alternation task, the rats experience multiple days and sessions where they get rewarded at any arm that they visit (provided it is not an immediate repeat). During this period of the behavior, the duration of a session is defined by a fixed number of rewards, or a fixed amount of time on the track (15 minutes), whichever came first. During the alternation task the duration of a session was defined either by a fixed number of center arm visits and at least one subsequent visit to any other arm, or a fixed amount of time on the track (15 minutes), whichever came first.

The algorithm underlying the spatial alternation task is such that three arms on the track have the potential for reward: arms 2, 3, and 4 have the potential to be rewarded, and arms 1, 5, and 6 do not. Of those three potentially rewarded arms we refer to the middle of the three arms as the center arm (arm 3) and the other two arms as the outer arms (arms 2 and 4). Reward is delivered at the center arms if and only if: 1) the immediately preceding arm whose reward well infrared beam was broken was not the center arm. Reward was delivered at the outer two arms if and only if: 1) the immediately preceding arm whose reward well infrared beam was broken was the center arm, and 2) prior to breaking the infrared beam at the center arm, the most recently broken outer arm infrared beam was not the currently broken outer arm infrared beam. The one exception to the outer arm rules was at the beginning of a session, if no outer arm infrared beam was broken prior to the first infrared beam break at the center arm, then only the first condition had to be met.

For the running of the behavior, the infrared beam break determined an arm visit; however, the rats sometimes go down an arm, get very close to the reward wells, but do not break the infrared beam. Therefore, for all the analyses described for the rats, an arm choice is defined as when a rat gets close to a reward well. These times were extracted from a video recording of the behavior. This proximity-based definition of an arm visit added additional arm visits to those defined by the infrared beam breaks, and none of them could ever be rewarded, nor alter the logic of the underlying algorithm. However, because of the non-Markovian nature of the reward contingency, they could affect the rewards provided for subsequent choices.

For the initial dataset, a total of 46 rats that underwent control surgery (22 males, 24 females) and 45 rats that underwent hippocampal lesion surgery (23 males, 22 females) were run on the spatial alternation task. 6 of the rats that underwent hippocampal lesion surgery showed no evidence of lesioning on CT and were therefore not included in the analysis. For the replication dataset, a total of 9 rats that underwent control surgery (all males) and 11 rats that underwent hippocampal lesion surgery (all males) were run on the spatial alternation task. For the replication dataset, all of the rats that underwent hippocampal lesion surgery showed evidence of lesioning on CT. Rats were housed together prior to their undergoing surgery. They were single housed for post-operative recovery and through food restriction and the behavior.

The first 800 choices during the spatial alternation behavior for each rat were used. Only 1 rats did not reach the criterion. That control, female, rat from the initial dataset, completed 618 choices, and was included in the analysis.

## Data and Code availability

Rat spatial alternation data generated for this work and code used to calculate CBAS will be posted to Github upon publication.

## Acknowledgements

We thank Claire Gillan and Benjamin de Bivort for making their data publicly available. We thank Thomas Akam, Nathanial Daw, and Adam Frank for very helpful comments on the manuscript. This research used resources of the Advanced Light Source, a DOE Office of Science User Facility under contract no. DE-AC02-05CH11231.

## Funding

This work was supported by grants from the Jane Coffin Childs Memorial Fund for Medical Research (D.B.K.), the UCSF Physician Scientist Scholars Program (D.B.K.), an NIH R25 (R25MH060482) (D.B.K), a Simons Foundation Autism Research Initiative grant (899599) (D.B.K.), an NIMH K08 Award (K08MH139983) (D.B.K.), the Max Planck Society (P.D.) and the Humboldt Foundation (P.D.).

## Author Contributions

D.B.K. and P.D. designed the study, D.B.K., J.P.R., and P.D. developed the analysis method, D.B.K., N.O.Y., C.Y.L., C.H., J.T., A.C., M.M., V.K. and G.W. collected the data, D.L.P. provided expertise and resources. D.B.K., N.O.Y., C.Y.L., and M.M. analyzed the data, D.B.K and P.D. wrote the manuscript, and D.B.K., J.P.R. and P.D. edited the manuscript.

## Declaration of interests

The authors declare no competing interests.

## Figure Legends

**Supplementary Figure 1.**
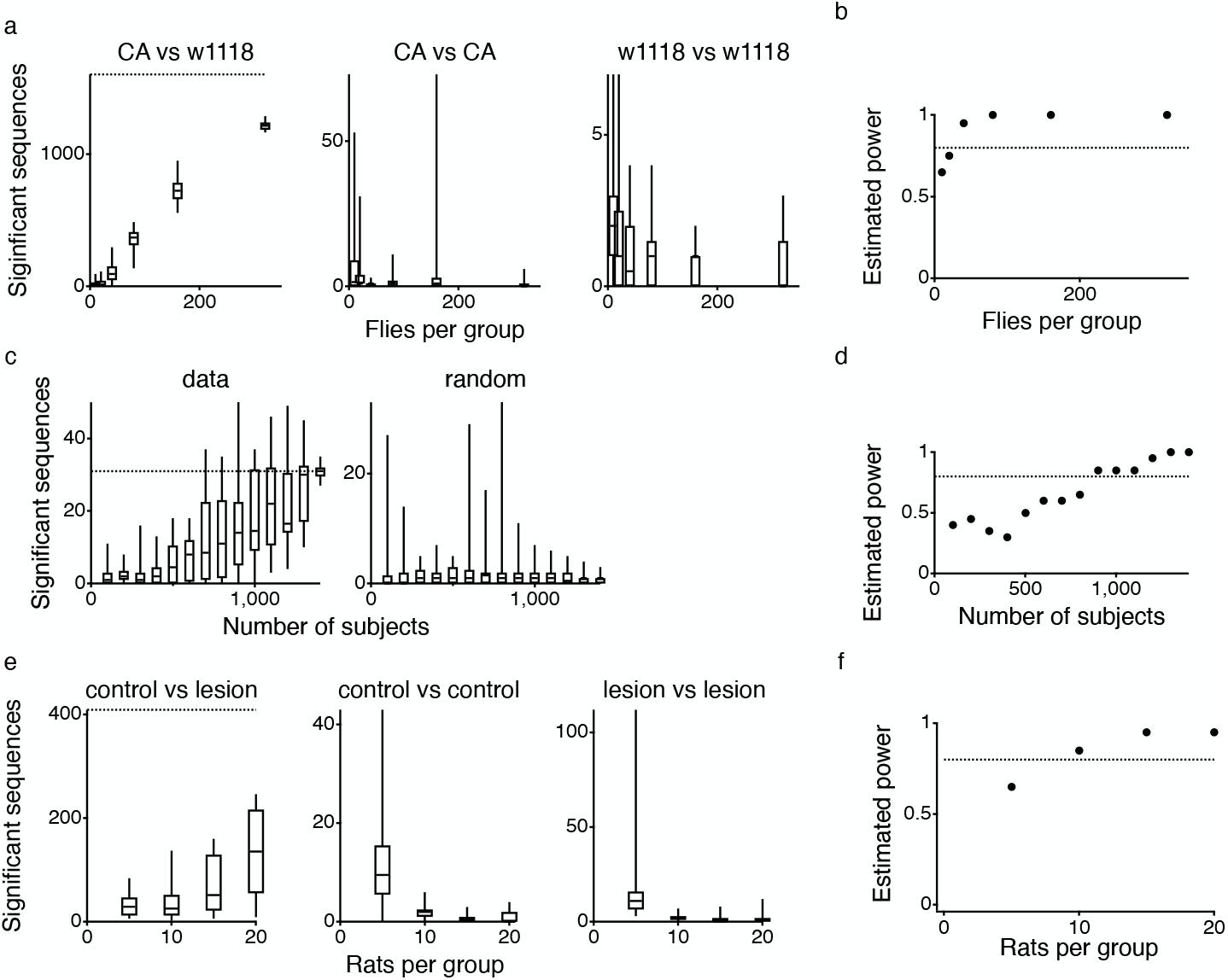
Sample size calculation for CBAS. (**a**) Flies: Left: the number of significant sequences when randomly resampling the populations without replacement and calculating CBAS on the smaller sample sizes. Middle: the number of significant sequences when comparing smaller samples sizes of the CA strain to itself, with nonoverlapping individuals in each group. Right: the number of significant sequences when comparing smaller samples sizes of the w1118 strain to itself, with nonoverlapping individuals in each group. (**b**) Power estimate for different sized groups of flies in each strain (see methods). (**c**) Humans: Left: the number of sequences significantly correlated with CBIT when randomly resampling the population separately without replacement and calculating CBAS on the smaller sample sizes. Right: the number of significant sequences when randomly resampling the population without replacement and comparing it to randomly generated CBIT scores drawn from a distribution imputed from the original CBIT scores and calculating CBAS on the smaller sample sizes. (**d**) Power estimate for different sized groups of human subjects (see methods). (**e**) Rats: Left: the number of significant sequences when randomly resampling the populations without replacement and calculating CBAS on the smaller sample sizes. Middle: the number of significant sequences when comparing smaller samples sizes of the control rats to itself, with nonoverlapping individuals in each group. Right: the number of significant sequences when comparing smaller samples sizes of the hippocampal lesion rats to itself, with nonoverlapping individuals in each group. (**f**) Power estimate for different sized groups of rats (see methods). For **a, c**, and **e** data points are overlayed by box and whisker plots. The center line of the box displays the median of the data, the top and bottom lines of the box show the 25^th^ and 75^th^ quartiles, respectively, and the end of the whiskers show the full range of the data. Additionally, the horizontal dotted lines for the left-most plot indicates the total number of significant sequences CBAS find in the entire dataset. For **b, d**, and **f** horizontal dotted line shows a value of 80%.

**Supplementary Figure 2.**
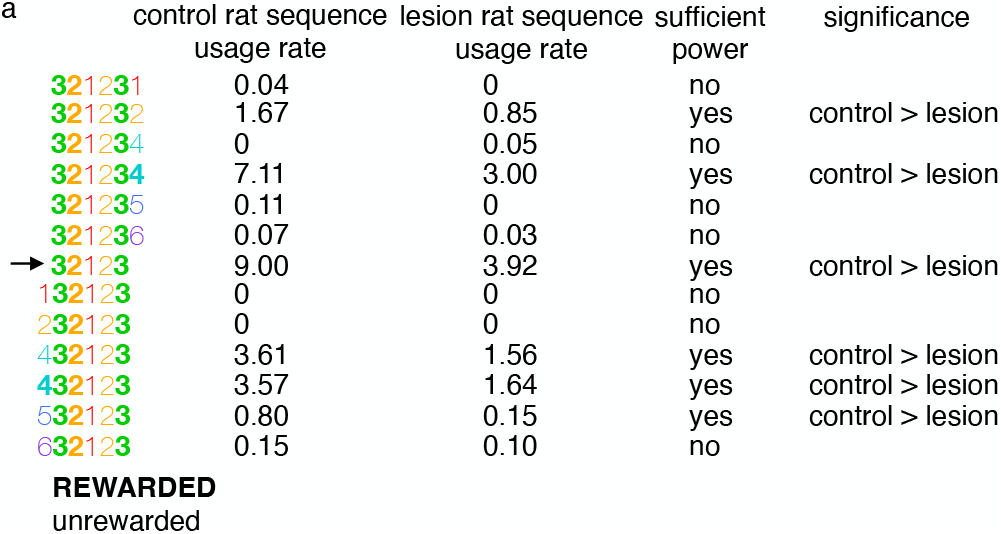
“Complete” sequence calculation. (**a**) A sequence unit is evaluated, and all sequences in the dataset that contain that unit are found. For the example shown, the sequence unit is: **32**12**3** (indicated by an arrow). As usual, emboldened numbers indicate that that choice was rewarded. Sequences are colored as in Fig 5 and 6. The second column lists the rate at which each sequence was utilized by control animals (as this sequence occurs more in control than lesion rats). The rate in the control population above which there is 80% power to significantly detect a sequence difference was determined to be 0.174. The third column lists whether sequences occurred above the power level. The fourth column indicates which sequences were determined to be significant, and in what direction. A “complete” sequence is therefore defined as a sequence for which all longer sequences that contain that sequence of interest, and which occur above a rate that allows for 80% power for differentiation, are found by CBAS to differ significantly in the same direction as the sequence of interest (e.g., are all found to be more prevalent in the control, than the lesion, rats).

**Supplementary Figure 3.**
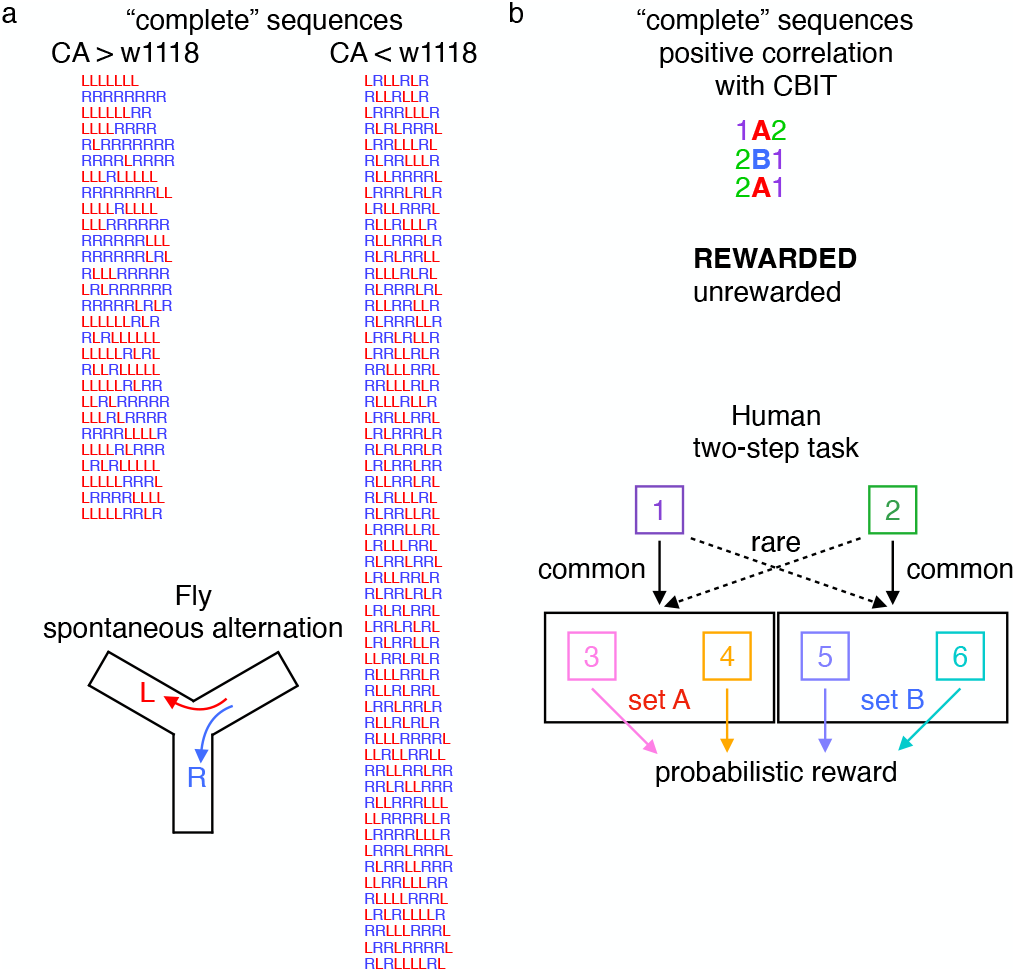
“Complete” sequences for fly and human tasks. All the sequences found to be complete for the fly task (**a**) and the human task (**b**) are shown. There is no “complete” sequence that is negatively correlated with CBIT for the human two-step task. For **b** bold and upper-case letters indicate that reward was received for that choice.

**Supplementary Figure 4.**
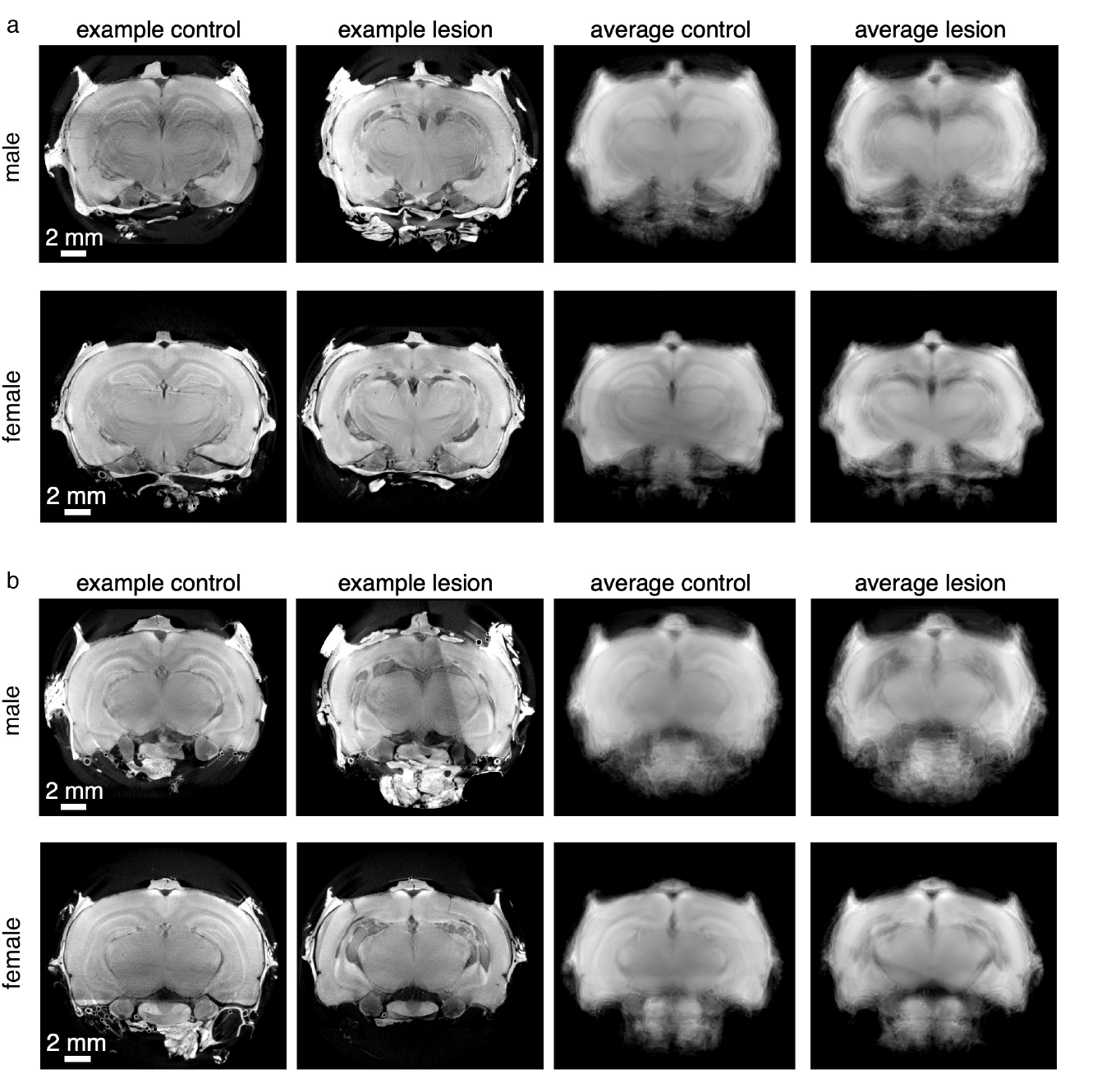
Micro-CT used to confirm hippocampal lesioning. Example virtual slices, displaying two different parts of the hippocampus (**a**, anterior, and **b**, posterior), from the micro CTs of a control and lesioned brain for males (top) and females (bottom). Additionally, the averages for the virtual slices for the control males (n = 22) and females (n = 18) and lesioned males (n = 24) and females (n = 21) are displayed. The blur in the averaging is due to imperfect registration across the different brains, as only a linear transformation was used for the registration. Note that hippocampal cell layers remain well defined and intact in the average images for the control, but largely absent in the lesioned average images, as revealed by the irregular darker areas with no tissue in the location of the hippocampus. The scale of the images are the same between example control and lesion images, and between average control and lesion images, but the scale of the average images have an increased gain compared to the example images to better highlight the features in the average images.

## Notes

### Competing Interest Statement

The authors have declared no competing interest.

### Summary of Updates

The entire manuscript was edited and updated.

